# A comparison between Mini-loop mediated isothermal amplification and polymerase spiral reaction for selective amplification of short template DNA

**DOI:** 10.1101/2022.05.19.492708

**Authors:** RC Allsopp, G Alexandrou, C Toumazou, S Ali, Coombes R Charles, M Kalofonou, J A Shaw

**Author notes:** **Corresponding author:** Rebecca Allsopp.

## Abstract

Isothermal amplification of circulating tumour-derived DNA (ctDNA) in the blood plasma could provide a rapid and cost effective alternative to PCR and NGS approaches for real-time cancer monitoring. Several variations of isothermal technologies exist, typically designed over unconstrained template length. Here, we compared the amplification efficiency of a compact loop mediated isothermal amplification reaction (termed AS-Mini-LAMP) with polymerase spiral reaction (PSR) suitable for analysis of ctDNA. Utilising 4-primers and targeting a 155 bp template encompassing the estrogen receptor (*ESR1)* c.1138G>C (p.E380Q) missense mutation.

Using synthetic E380Q template DNA and Bst2.0 polymerase, results demonstrate that AS-Mini-LAMP was capable of selective mutant allele DNA amplification to a limit of 1,000 mutant copies, whereas no specific amplification was observed by PSR. The alternative use of Bst3.0 polymerase for either AS-Mini-LAMP or PSR revealed non-canonical events that underpin potentially misleading results when employing isothermal chemistries. In conclusion, AS-Mini-LAMP is more suited to mutation detection than PSR.

## Introduction

Cancer is one of the leading causes of death worldwide with tissue biopsy remaining as the current gold standard for diagnosis. However, single biopsy specimens of primary tumours are unlikely to fully represent the genetic diversity of malignancy or contain sufficient number of events for next generation sequencing (NGS) based detection. Furthermore, anticancer treatments may cause additional selection pressures and influence the mutational nature within a tumour (1). Clinical practice therefore requires regular assessment of complementary molecular biomarkers such as present within minimally invasive liquid biopsy in order to monitor and molecularly profile potentially important changes during disease evolution.

NGS monitoring of plasma derived circulating tumour DNA (ctDNA) have demonstrated proof of principle utility (2-5). However, isothermal amplification reactions present attractive, low cost and rapid alternatives, requiring no expensive equipment or informatics expertise. Our previous work, utilising variant selective loop-mediated isothermal amplification with additional unmodified self-stabilising competitive primers to further reduce non-specific amplification (termed USS sbLAMP) (6) demonstrated selective and rapid amplification of *PIK3CA* c.3140A > G p.H1047R in tumour tissue DNA (7) and *ESR1* mutations c.1138G>C p.E380Q + c.1610A>C p.Y537S in plasma cell free DNA (cfDNA) (8). Furthermore, these reactions were successfully piloted on an ISFET (ion-sensitive field-effect transistor) based CMOS integrated Lab-on-Chip (LoC) system, demonstrating efficacy of use on a miniaturised, portable and scalable interface.

Of challenge, classic LAMP reactions (9) and our own USS-sbLAMP reactions detailed above (7, 8), typically span greater than 200 bp of template DNA, outside the range of cfDNA (∼ 167 bp) and ctDNA (< 167 bp) fragments (10-13). To overcome this, we developed a more compact allele selective LAMP plus USS reaction termed AS-Mini-LAMP + USS (14) spanning only 155 bp of template DNA, within the range of cfDNA and ctDNA. However, in each of these cases (classic LAMP, USS-sbLAMP and AS-Mini-LAMP + USS) the number of primers required (6, 8 and 6 respectively) opens up the possibility of non-specific amplification, a key challenge for any LAMP optimisation (15). This study therefore investigates the possibility of a less complex 4-primer approach, also easier to compact over short cfDNA-like template. AS-Mini-LAMP (absence of additional USS primers) is compared to an alternative isothermal amplification strategy termed polymerase spiral reaction (PSR) (16), which to date has only been demonstrated for the detection of infectious diseases where reaction template span is not of concern (17, 18). Targeting identical template annealing sequences surrounding the *ESR1* E380Q missense mutation (and target of our previous investigations for comparison (8, 14)), this study directly compares the ability of AS-Mini-LAMP and PSR to amplify across a short stretch of synthetic DNA, optimal for cfDNA workflows.

Supplementary Figure 1A demonstrates this 4-primer approach. The forward and reverse inner primers target template DNA sequence at their 3’-end whilst encoding unique modes of secondary exponential amplification at their 5’-end. For AS-Mini-LAMP (Supplementary Figure 1B) stem-loop formation (through self-hybridisation of the 5’-FIP linked F1c sequence to self-complementary F1 template region in the newly synthesized strand) is encoded at the 5’-end. For PSR (Supplementary Figure 1C), product spiral formation (via hybridisation of primer introduced reverse complementary sequences of botanic origin) is encoded at the 5’-end. A second primer pair, common to both AS-Mini-LAMP and PSR serve to facilitate strand displacement (Supplementary Figure 1A, B, C). Both strategies employ Bst-like polymerase, performing optimally at ∼ 65°C with inherent strand displacement activity and inhibitor tolerance, thus minimising the need (upon optimisation) for sample purification (19, 20). In this study, we additionally compared the use of the Bst2.0 *in silico* homolog of Bst DNA polymerase large fragment with the further improved Bst3.0 polymerase.

## Methods

### DNA template preparation

A 402bp double-stranded synthetic DNA sequence containing the *ESR1* c.1138G>C p.E380Q missense mutation (Cosmic ID COSM3829320) was synthesised by Integrated DNA Technologies. DNA was resuspended in nuclease free water to a concentration of 10 ng/μL aliquoted to single use samples and stored at −20 °C. Template DNA was diluted to 1×10^4^ copies per isothermal reaction.

### Isothermal assay design

AS-Mini-LAMP and PSR assays, targeting identical *ESR1* E380Q surrounding regions were designed to span a minimal template sequence using a 4-primer strategy. Primers were analysed using the online Oligo Analyser Tool in the IDT database (http://eu.idtdna.com/calc/analyzer) to identify thermodynamically favourable combinations with minimal self-complementarity or hairpin-loop structures. Chosen primer design were HPLC purified. Note that AS-Mini-LAMP but not PSR encodes selectivity for the E380Q mutation at this stage of comparative short template amplification (Supplementary Figure 2 and 3 respectively).

### Isothermal reactions

Bst2.0 enzyme (isothermal buffer pack) and Bst3.0 enzyme (isothermal buffer II) were compared. All reaction were prepared to a final volume of 15 ul, performed at a constant temperature of 65 °C and monitored for 80 minutes in a StepOne Real-Time PCR System (Applied Biosystems) measuring EvaGreen dye intercalation. Unless otherwise started, reagents were purchased from New England BioLabs. Experiments were prepared in triplicate and performed twice. Each reaction consisted of:

### LAMP

1 x enzyme specific isothermal buffer, 8 mM MgSO_4_, 1.4 mM dNTPs mix, 10 x primer master mix (1.6 µM FIP / BIP, 0.2 µM F3 / B3) (Sigma, HPLC purified), 1 x EvaGreen dye (Biotium), 3 µl DNA template dilution (1×10^4^ copies/reaction) and 0.32 U/µl Bst enzyme made up to a final reaction volume of 15μl with nuclease-free water.

### PSR

1 x enzyme specific isothermal buffer, 8 mM MgSO_4_, 1.4 mM dNTPs mix, 10 x primer master mix (1.6 µM FP / RP, 0.8 µM IF / IB) (Sigma, HPLC purifed), 1 x EvaGreen dye (Biotium), 3 µl DNA template dilution (1×10^4^ copies/reaction) and 0.32 U/µl Bst enzyme made up to a final reaction volume of 15μl with nuclease-free water.

### Data analysis and statistics

Amplification efficiency was quantified by the separation in ‘time to positive’ (TTP) between target template amplification and non-specific no template control (NTC) amplification exceeding the manually set signal threshold (0.2; point at which amplification exceeds and enters linear, maximum amplification rate), determined in a continuously monitored reaction. Data are presented as mean TTP ± S.E.M and are an average of 2 independent experiments each performed in triplicate. Visualisation of the characteristic amplicon ladder pattern consistent with spiral formation was by gel electrophoresis on 2% agarose gel imaged using a BioRad GelDoc XR+System transilluminator. Reaction efficiency is calculated by E=-1+10^(−1/slope)^. Statistically significant differences were calculated by one-way ANOVA followed by Dunnett’s multiple comparisons test (GraphPad Prism 8.1.2).

### Agarose gel electrophoresis and restriction digest confirmation

Isothermal products (1 μL) were digested with *AvaII* (New Englanc Biolabs) according to manufacturer’s instructions to confirm template specific origin (Supplementary figure 1). Calculation of the expected banding pattern of LAMP products is complex but supported through visual representation on a virtual gel using the online tool http://creisle.github.io/creisle.lamprflp/index.html. Calculation of the expected banding pattern of PSR products is simple, a single restriction digest site resulting in spiral digestion to a single sized easily calculable product.

## Results

Parallel AS-Mini LAMP and PSR investigations targeting 155 bp of template DNA, surrounding the *ESR1* c.1138G>C p.E380Q mutation, were performed independently using Bst2.0 or Bst3.0 polymerase. In the case of Bst2.0 LAMP (Figure 1A), mutant template amplification occurred at 53.5 ± 1.4 min with no amplification present within the NTC. In contrast, Bst3.0 LAMP amplification of both mutant template and NTC occurred in unison and significantly earlier at 24.2 ± 0.4 and 28.7 ± 1.6 (Figure 1B). In attempt to understand these Bst induced differences, FIP only and BIP only LAMP reactions were prepared using Bst2.0 (Figure 1C and 1E respectively) or Bst3.0 (Figure 1D and 1F respectively). Bst2.0 LAMP FIP only and BIP only reactions demonstrated no amplification. In contrast, Bst3.0 LAMP FIP only and BIP only reactions resulted in overlapping non-specific mutant template and NTC amplification at 28.8 ± 0.7 and 30.6 ± 0.4 min respectively for FIP only and 37.5 ± 2.1 and 38.7 ± 1.1 min respectively for BIP only. Outer primers to assist strand displacement were included in all reactions.

**Figure 1.**
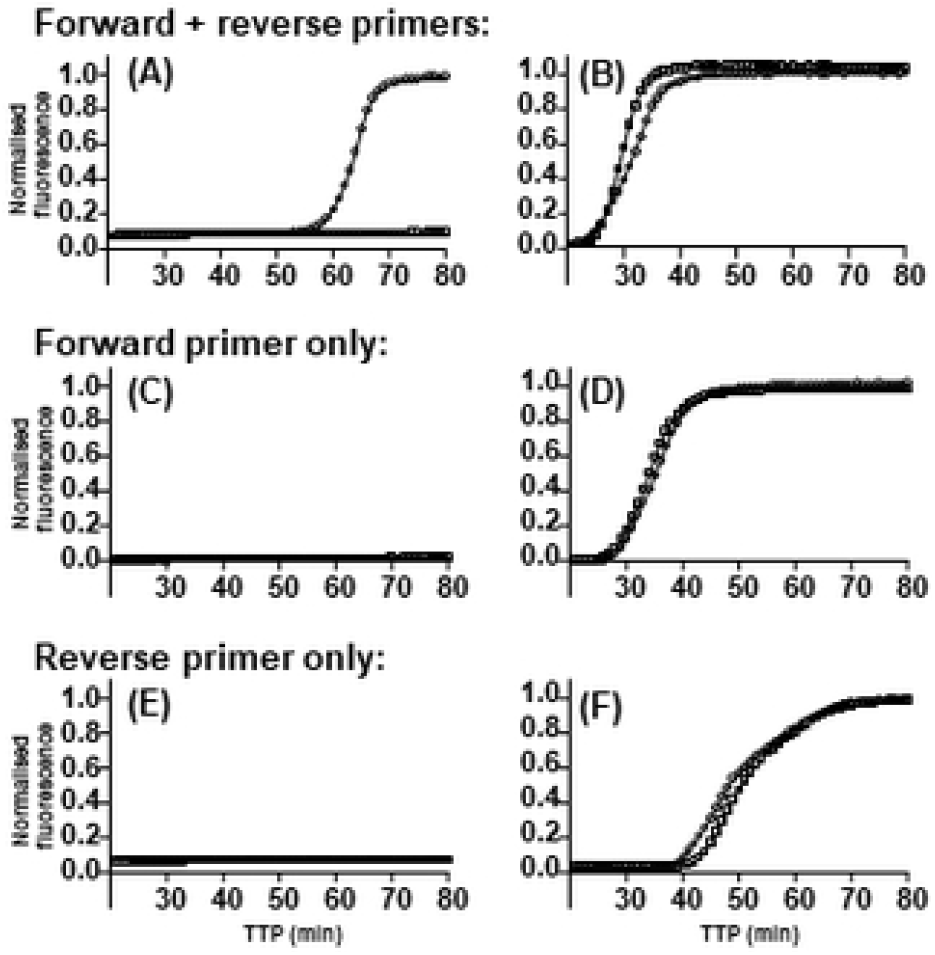
Influence of Bst polymerase on LAMP. Representative LAMP amplification using (A+B) forward and reverse reaction primers, (C+D) forward primer only and (E+F) reverse primer only in the presence of either Bst2.0 (A, C and E) of Bst3.0 polymerase (B, D and F). E380Q template reactions contain 1×10^4^ copies of template DNA (◯) or NTC (□). Reactions were performed at 65°C for 80 minutes monitored continuously by EvaGreen incorporation. Time to positive (TTP) is the time (minutes) at which amplification exceeds the manually set, reaction consistent threshold (red dotted line) and enters the rapid linear, exponential phase. Data representative of two independent experiments each performed in triplicate.

In the case of PSR (Figure 2), complete forward primer (FP) and reverse primer (RP) Bst2.0 mutant template and NTC reactions (Figure 2A) resulted in no amplification, whereas FP+RP Bst3.0 reactions (Figure 2B) resulted in non-specific simultaneous mutant template and NTC amplification at 43.6 ± 2.0 and 43.4 ± 1.7 min respectively. Bst2.0 FP or RP only reactions (Figure 2C and 2E respectively) resulted in no amplification. In contrast, Bst3.0 initiated FP or RP only reactions (Figure 2D and 2F respectively) resulted in overlapping mutant template and NTC amplification at 34.6 ± 1.6 and 38.2 ± 0.7 min for FP only and 58.4 ± 2.5 and 55.0 ± 0.96 min RP only. Outer primers to assist strand displacement were included in all reactions.

**Figure 2.**
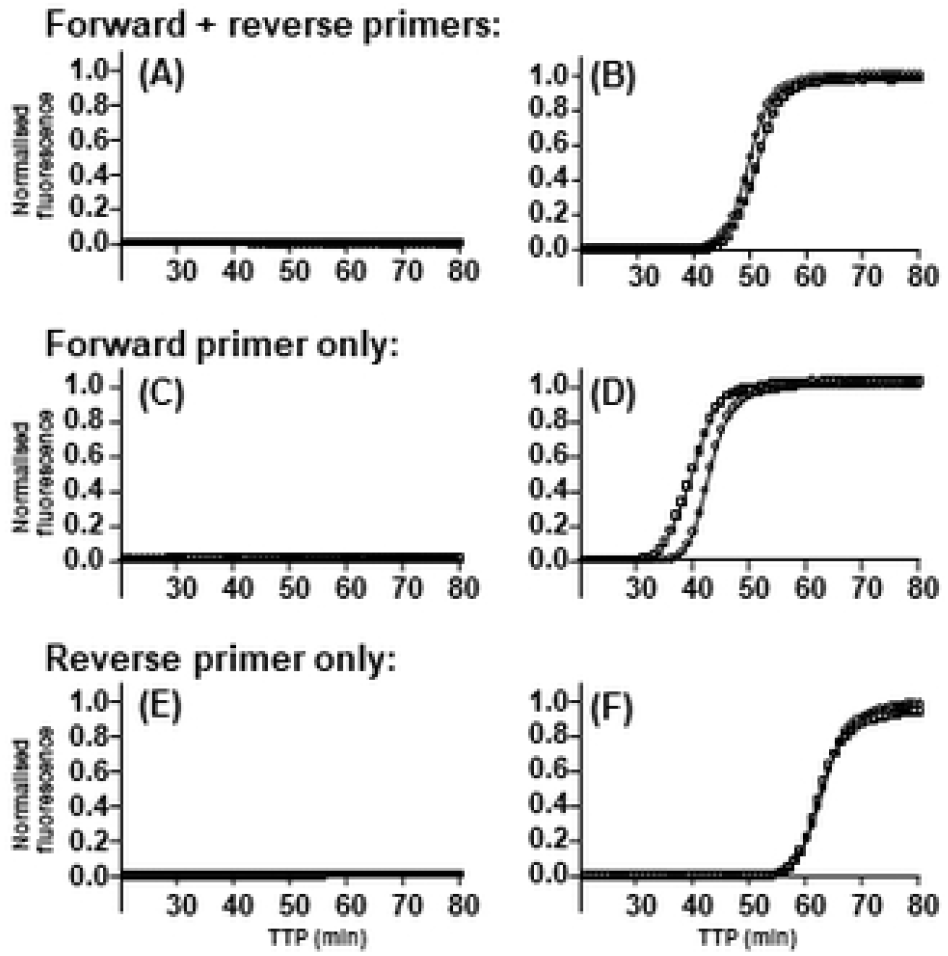
Influence of Bst polymerase on PSR. Representative PSR amplification using (A+B) forward and reverse reaction primers, (C+D) forward primer only and (E+F) reverse primer only in the presence of either Bst2.0 (A, C and E) of Bst3.0 polymerase (B, D and F). E380Q template reactions contain 1×10^4^ copies of template DNA (◯) or NTC (□). All reactions were performed at 65°C for 80 minutes monitored continuously by EvaGreen incorporation. Time to positive (TTP) is the time (minutes) at which amplification exceeds the manually set, reaction consistent threshold (red dotted line) and enters the rapid linear, exponential phase. Data representative of two independent experiments each performed in triplicate.

Bst2.0 AS-Mini-LAMP (FIP+BIP) was the only reaction resulting in significant delay (26.5 minutes) between selective mutant template and non-selective NTC amplification. The assay limit of detection and selectivity of mutant allele over wild type (WT) were determined using independent ten-fold serial template dilutions (1×10^7^ to 1×10^3^) of mutant and WT template synthetic DNA (Figure 3A and B respectively); linear regression analysis demonstrating a higher reaction efficiency of *E380Q* AS-Mini-LAMP for mutant template DNA (R^2^ 0.9524) over WT template DNA (R^2^ 0.8175). Moreover, serial dilutions showed AS-Mini-LAMP selectively amplified mutant allele > 10 minutes ahead of WT allele to a limit of 1,000 template copies (Supplementary Table 1). As confirmation, *AvaII* restriction enzyme digestion of AS-Mini-LAMP products (Figure 3C) demonstrated the predicted fragment sizes (90 bp, 75 bp, 60 bp and 10 bp). Lastly, the ability of Bst2.0 AS-Mini-LAMP to detect mutant E380Q target within a background of WT allele was also investigated, as a surrogate for ctDNA detection within total cfDNA background. Totalling 1×10^4^ copies, decreasing ratio of mutant : WT allele were tested including 100:0, 75:25, 50:50, 25:75 and 0:100. Results (Figure 3D and Supplementary Table 2) demonstrated that AS-Mini-LAMP was able to selectively amplify mutant target template >10 minutes ahead of WT for all mixed population ratios tested.

**Figure 3.**
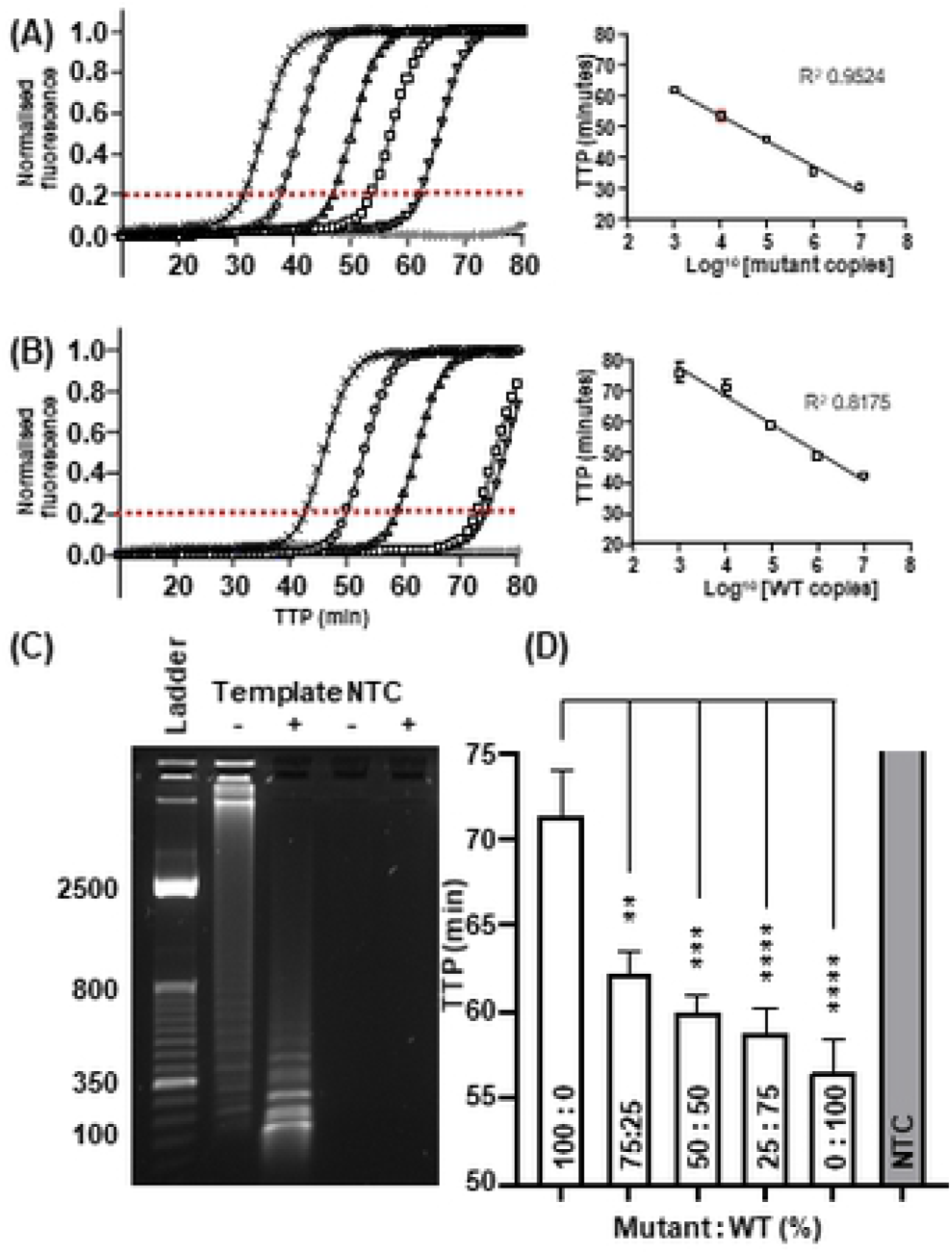
Mutant selective AS-Mini-LAMP amplification of *ESR1* E380Q template DNA in a mixed WT population, performed using Bst2.0 polymerase. Representative AS-Mini-LAMP amplification of serially diluted synthetic (A) E380Q mutant and (B) WT template DNA including 1×10^7^ (x), 1×10^6^ (□), 1×10^5^ (△), 1×10^4^ (□), 1×10^3^ (▽), NTC shown in grey (*). Linear regression analysis (R^2^ value) demonstrates the reaction efficiency against selective mutant and non-selective WT template. C) Bst2.0 AS-Mini-LAMP reaction product pre and post (−/+) restriction enzyme digestion using *AvaII* at 37°C for 15 minutes. Lane M is the molecular weight marker. Undigested template products were visualised at high molecular weight compared to the ladder pattern concurrent with predicted sizes 90 bp, 75 bp, 60 bp and 10 bp upon digestion (D) AS-Mini-LAMP mutant allele selective amplification in a mixed mutant : WT population. Time to positive (TTP) is the time (minutes) at which amplification exceeds the manually set, reaction consistent threshold (red dotted line) and enters the rapid linear, exponential phase. Data are shown as the mean ± S.E.M, representative of two independent experiments each performed in triplicate, \*\**P* = 0.0027; \*\*\**P* = 0.001; \*\*\*\**P* < 0.0001.

## Discussion

Herein we directly compared the ability of two 4-primer design isothermal amplification strategies, AS-Mini-LAMP and PSR, to amplify a short (155 bp) region of synthetic template DNA as a surrogate for ctDNA. The choice and effect of strand displacement polymerase enzyme was also investigated. Targeting identical template sequences, Bst2.0 AS-Mini-LAMP but not Bst2.0 PSR demonstrated selective *ESR1* E380Q mutant allele detection, to a limit of 1,000 copies and > 10 minutes ahead of non-selective WT allele amplification within a mixed allele reaction. Restriction enzyme digestion confirmed amplification products originated from input template DNA. In contrast, Bst2.0 PSR failed to amplify mutant template DNA at equivalent and significantly higher number of copies of mutant allele DNA (1×10^4^ to 1×10^7^). Whilst the purpose of this paper is to compare the ability of Mini-LAMP and PSR at identical template annealing locations, alternative thermodynamically favourable PSR primers targeting nearby regions were also tested, equally failing to amplify (data not shown). AS-Mini-LAMP and PSR differ significantly in their mode of secondary exponential amplification as determined by the sequence encoded at the 5’-end of the inner forward and reverse primer pair. Within AS-Mini-LAMP, formation of a stem-loop intermediary structure leads to extension and formation of new sequence from the 3’-open end of the loop towards its 5’ end in a classical anti-parallel direction (21). In contrast, when a displaced complete forward and reverse reaction PSR product strand self-anneals, extension and new product formation proceeds from the 3’-open end, spiralling in a parallel direction. In nature, nucleic acids can adopt a range of confirmations, and whilst a parallel strand orientation is very stable, it has been shown to be less stable than its antiparallel counterpart (22). Thus, raising the suggestion that exponential amplification of a LAMP intermediary stem-loop structure in an anti-parallel direction may offer enhanced efficiency due to added structural stability compared to the spiral amplification of a PSR product in a parallel direction.

Performing the same reactions using Bst3.0 in place of Bst2.0 demonstrated non-specific miss-annealing within these long length primer reactions (43 bases). For both AS-Mini-LAMP and PSR, use of Bst3.0 polymerase resulted in rapid, overlapping non-specific mutant template and NTC amplification, evident in single forward of reverse inner primer reactions, thus confirming the non-specific nature of these amplification events. Most likely, this can be ascribed to Bst3.0 polymerase’s enhanced reverse transcriptase activity and promotion of miss-annealing events. A similar observation identified multimeric DNA production in reactions using a linear template, one primer, and strand-displacement polymerase with RT activity, termed ‘unusual isothermal multimerization and amplification’ or UIMA, a phenomena rarely seen when utilizing polymerases without RT activities (23). Similar events have also been reported using Bst2.0 polymerase. However, optimisation of reaction conditions can promote efficient specific isothermal amplification. For instance, the presence of 0.25 fold SYBR Green intercalating dye inhibited multimerization in Bst2.0 reactions, whereas over 1.25 fold was required for multimerization inhibition in Bst 3.0 polymerase reactions (24). Our finding provide further evidence of such non-specific isothermal amplification events and reinforce the importance of additional secondary product confirmation measures (restriction digest or melt curve analysis).

Within the context of our previous efforts, it is clear that the simplified 4-primer AS-Mini-LAMP (demonstrating sensitivity of 1,000 mutant copies in ∼ 60 min and > 10 minutes ahead of non-selective WT amplification) or PSR approach (no amplification) offered no advantages over our previously described 8 and 6 primer reactions (8, 14). False positives (the result of non-specific amplification) remained a challenge, although largely driven by polymerase choice in this case. Moreover, a significant sacrifice in amplification velocity resulted. Our equivalent 6-primer ‘AS-Mini-LAMP + USS’ alterative assay, also spanning 155 bp, provided 2-fold enhanced reaction sensitivity and velocity (500 mutant copies in ∼ 25 minutes) and greater allele selectivity (> 20 minutes ahead of non-selective WT amplification) (14). However, it is important to note that these AS-Mini-LAMP assays target overlapping but subtlety different (a few base pairs) template annealing locations surrounding E380Q, so primer thermodynamic properties may factor here. Finally, our 8-primer E380Q ‘USS sbLAMP’ assay designed over unconstrained template length (∼200 bb) also maintained ability to selectively amplify fragmented template DNA. Fragments of 90 bp, created by restriction digest (maintaining only the F1c and B1c binding sites) amplified only 6 minutes later than undigested template DNA (∼ 25 minutes) to a limit of 1,000 copies and > 10 minutes ahead of non-selective WT template amplification (8).

In summary, selective *ESR1* E380Q detection within short template DNA is possible using either USS-sbLAMP (designed over ∼200 bp template DNA) (8) or the more compact AS-Mini-LAMP + USS design strategy (155 bp) (14). Amplification performed across shorter templates are likely to be less limited by strand displacement polymerase activity compared to reactions performed on longer templates (9), possibly underpinning the reason behind AS-Mini-LAMPs enhanced limit of detection. In contrast, PSR is deemed more suitable for the detection of pathogens across unconstrained template length rather than short template mutation analysis.

## Data availability

All data generated or analysed during this study are included in this published article. Data generated and analysed throughout the study can be made available from the corresponding author on reasonable request.

## References

1. Fisher R, Pusztai L, Swanton C. Cancer heterogeneity: implications for targeted therapeutics. Br J Cancer. 2013;108(3):479–85.

2. Guttery DS, Page K, Hills A, Woodley L, Marchese SD, Rghebi B, et al. Noninvasive detection of activating estrogen receptor 1 (ESR1) mutations in estrogen receptor-positive metastatic breast cancer. Clin Chem. 2015;61(7):974–82.

3. Klein EA, Hubbell E, Maddala T, Aravanis A, Beausang JF, Filippova D, et al. Development of a comprehensive cell-free DNA (cfDNA) assay for early detection of multiple tumor types: The Circulating Cell-free Genome Atlas (CCGA) study. Journal of Clinical Oncology. 2018;36(15_suppl):12021-.

4. Kodahl AR, Ehmsen S, Pallisgaard N, Jylling AMB, Jensen JD, Laenkholm AV, et al. Correlation between circulating cell-free PIK3CA tumor DNA levels and treatment response in patients with PIK3CA-mutated metastatic breast cancer. Mol Oncol. 2018;12(6):925–35.

5. Page K, Guttery DS, Fernandez-Garcia D, Hills A, Hastings RK, Luo J, et al. Next Generation Sequencing of Circulating Cell-Free DNA for Evaluating Mutations and Gene Amplification in Metastatic Breast Cancer. Clin Chem. 2017;63(2):532–41.

6. Malpartida-Cardenas K, Rodriguez-Manzano J, Yu L-S, Delves MJ, Nguon C, Chotivanich K, et al. Allele-Specific Isothermal Amplification Method Using Unmodified Self-Stabilizing Competitive Primers. Analytical Chemistry. 2018;90(20):11972–80.

7. Kalofonou M, Malpartida-Cardenas K, Alexandrou G, Rodriguez-Manzano J, Yu LS, Miscourides N, et al. A novel hotspot specific isothermal amplification method for detection of the common PIK3CA p.H1047R breast cancer mutation. Sci Rep. 2020;10(1):4553.

8. Alexandrou G, Moser N, Mantikas KT, Rodriguez-Manzano J, Ali S, Coombes RC, et al. Detection of Multiple Breast Cancer ESR1 Mutations on an ISFET Based Lab-on-Chip Platform. IEEE Trans Biomed Circuits Syst. 2021;15(3):380–9.

9. Notomi T, Okayama H, Masubuchi H, Yonekawa T, Watanabe K, Amino N, et al. Loop-mediated isothermal amplification of DNA. Nucleic Acids Res. 2000;28(12):E63–E.

10. Jiang P, Chan CWM, Chan KCA, Cheng SH, Wong J, Wong VW-S, et al. Lengthening and shortening of plasma DNA in hepatocellular carcinoma patients. Proceedings of the National Academy of Sciences. 2015;112(11):E1317–E25.

11. Liu HE, Vuppalapaty M, Wilkerson C, Renier C, Chiu M, Lemaire C, et al. Detection of EGFR Mutations in cfDNA and CTCs, and Comparison to Tumor Tissue in Non-Small-Cell-Lung-Cancer (NSCLC) Patients. Frontiers in Oncology. 2020;10.

12. Mouliere F, Robert B, Arnau Peyrotte E, Del Rio M, Ychou M, Molina F, et al. High fragmentation characterizes tumour-derived circulating DNA. PLoS One. 2011;6(9):e23418–e.

13. Underhill HR, Kitzman JO, Hellwig S, Welker NC, Daza R, Baker DN, et al. Fragment Length of Circulating Tumor DNA. PLoS Genet. 2016;12(7):e1006162–e.

14. Allsopp R, Alexandrou G, Toumazou C, Ali S, Coombes C, Kalofonou M, et al. Rapid Detection of Circulating Tumour DNA using Allele Specific Mini-Loop Mediated Isothermal Amplification 2022.

15. Varona M, Anderson JL. Advances in Mutation Detection Using Loop-Mediated Isothermal Amplification. ACS Omega. 2021;6(5):3463–9.

16. Liu W, Dong D, Yang Z, Zou D, Chen Z, Yuan J, et al. Polymerase Spiral Reaction (PSR): A novel isothermal nucleic acid amplification method. Scientific Reports. 2015;5(1):12723.

17. Sun W, Du Y, Li X, Du B. Rapid and Sensitive Detection of Hepatitis C Virus in Clinical Blood Samples Using Reverse Transcriptase Polymerase Spiral Reaction. J Microbiol Biotechnol. 2020;30(3):459–68.

18. Tran DH, Cuong HQ, Tran HT, L. UP, Do HDK, Bui LM, et al. A comparative study of isothermal nucleic acid amplification methods for SARS-CoV-2 detection at point of care. bioRxiv. 2020:2020.05.24.113423.

19. Jevtuševskaja J, Krõlov K, Tulp I, Langel Ü. The effect of main urine inhibitors on the activity of different DNA polymerases in loop-mediated isothermal amplification. Expert Rev Mol Diagn. 2017;17(4):403–10.

20. Oscorbin IP, Belousova EA, Boyarskikh UA, Zakabunin AI, Khrapov EA, Filipenko ML. Derivatives of Bst-like Gss-polymerase with improved processivity and inhibitor tolerance. Nucleic Acids Res. 2017;45(16):9595–610.

21. Watson JD, Crick FH. Molecular structure of nucleic acids: a structure for deoxyribose nucleic acid. J.D. Watson and F.H.C. Crick. Published in Nature, number 4356 April 25, 1953. Nature. 1974;248(5451):765.

22. Wunnicke D, Ding P, Yang H, Seela F, Steinhoff HJ. DNA with Parallel Strand Orientation: A Nanometer Distance Study with Spin Labels in the Watson-Crick and the Reverse Watson-Crick Double Helix. J Phys Chem B. 2015;119(43):13593–9.

23. Wang G, Ding X, Hu J, Wu W, Sun J, Mu Y. Unusual isothermal multimerization and amplification by the strand-displacing DNA polymerases with reverse transcription activities. Sci Rep. 2017;7(1):13928.

24. Garafutdinov RR, Gilvanov AR, Sakhabutdinova AR. The Influence of Reaction Conditions on DNA Multimerization During Isothermal Amplification with Bst exo-DNA Polymerase. Appl Biochem Biotechnol. 2020;190(2):758–71.

