## Supplementary material for "A comparison between Mini-loop mediated isothermal amplification and polymerase spiral reaction for selective amplification of short template DNA"

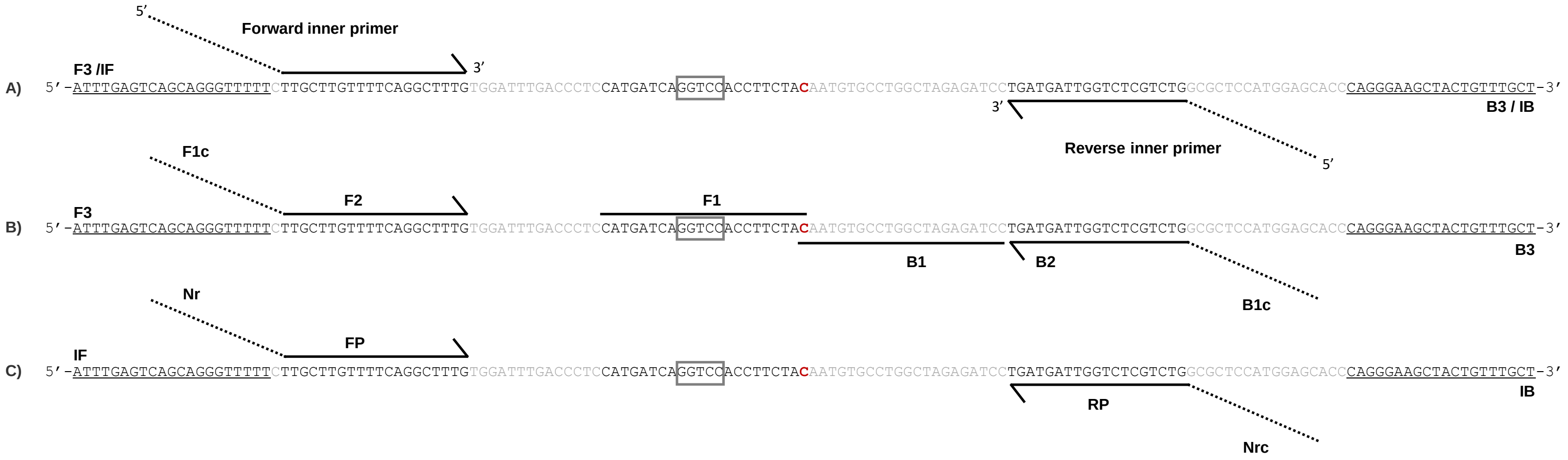


**Supplementary Figure 1. Allele selective mini- loop-mediated isothermal amplification (AS-Mini-LAMP) and polymerase spiral reaction (PSR) common *ESR1* template annealing primer sites. A)** The 3’-end of forward and reverse inner primers (solid line) target *ESR1* template regions (spanning 96 bp). Upon annealing and extension, product strand DNA is displaced by the action of outer sequence annealing primers termed F3/B3 or IF/IB in LAMP and PSR respectively (underlined and spanning 155 bp). **B) AS-Mini-LAMP reaction:** The 3’-F2/B2 regions (solid line) of the forward and reverse inner primer pair (FIP and BIP respectively) anneal and extend the *ESR1* template, incorporating 5’-F1c/B1c sequences (dotted line) of sequence complementarity (‘c’) to product strand regions F1 and B1, subsequent self-hybridisation results in the formation of LAMP stem-loop structures. Note that the variant nucleic acid responsible for the ESR1 E380 mutation (highlighted in red) is located at the terminal 5’-position of both F1c and B1c, conferring selective amplification over wild type allele. **C) PSR:** 3’-regions (solid line) of forward and reverse inner primers (FP, RP respectively) anneal and extend the *ESR1* template, incorporating reverse complementary (rc) 5’-sequences NR and Nrc (dotted line) of botanic origin, subsequent self-hybridisation results in PSR spiral formation. Note that this reaction is not E380Q selective but serves to comparatively assess an alternative isothermal amplification strategy targeting common sequence regions. The variant nucleotide base responsible for E380Q mutation is highlighted red (‘G’ within wild type sequence). The restriction digest site *AvaII* (boxed in grey) is utilised for diagnostic evaluation of isothermal amplification products.

**AS-LAMP Forward primer 'FIP' initiation:**

| Primer: | Sequence: |
|---|---|
| LAMP Mutant FIP | 5'-gtagaaggtggacctgatcatgTTGCTTGTTTTCAGGCTTTG-3' |
| LAMP Mutant BIP | 5'-caatgtgcctggctagagatccCAGACGAGACCAATCATCA-3' |
| F3 (IF) | ATTTGAGTCAGCAGGGTT |
| B3 (IB) | AGCAAACAGTAGCTTCCCTG |

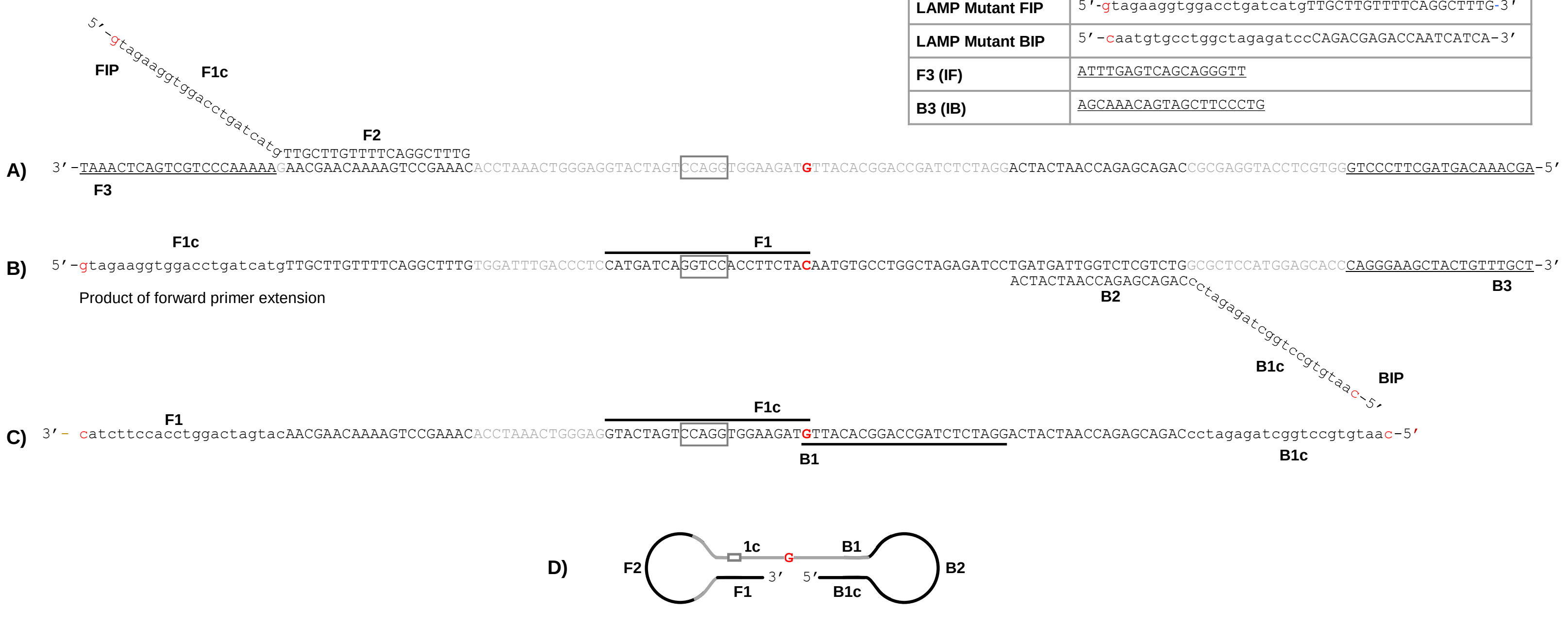


**Formation of a double-loop stem dumbbell structure**

Supplementary **Figure 2. Forward inner primer 'FIP' initiated ASS-Mini-LAMP reaction. A)** The 3'-F2 region of the inner FIP anneals and extends template DNA resulting in new strand synthesis, subsequently displaced by the action of outer F3 primer. Formation of a 5'-end loop structure can now occur due to sequence complementarity of the 5'-FIP linked F1c sequence to the F1 template region in the newly synthesized strand. **B)** Annealing and extension of the 3'-B2 region of the reverse BIP primer opens up this loop structure, forming a complementary strand which is again displaced by the action of the outer B3 primer. The resulting sequence (C) containing self- complementary regions encoding mutant allele selectivity (single base responsible for mutation shown in red) at both terminal 5'-positions is now able to form **(D)** a dumbbell shape, initiating self-primed extension from its open 3'-end. Simultaneously, the same reaction is occurring on the opposite single stranded template DNA initiated by reverse 'BIP' primer, resulting in self-primed extension from its open 3'-end with additional FIP/BIP annealing to these strands resulting in product of various lengths and multiple loop structures formed by annealing between alternatively inverted repeats of the target sequence in the same strand. These products appear as a ladder pattern by gel electrophoresis and restriction digest (using *AvaII*, grey boxed sequence) can be performed to confirm product identity. Restriction digest fragment expected size for LAMP reactions were calculated at 90 bp, 75 bp, 60 bp and 10 bp. Additional product specificity check (contamination control) is provided in the form of a melt curve of appropriate temperature.

**PSR Forward primer ‘FP’ initiation:**

| Primer: | Sequence: |
|---|---|
| **PSR FP** | 5’-acgattcgtacatagaagtatagTTGCTTGTTTTCAGGCTTTG-3’ |
| **PSR RP** | 5’-gatatgaagatacatgcttagcaCAGACGAGACCAATCATCA-3 |
| **IF (F3)** | ATTTGAGTCAGCAGGGTT |
| **IB (B3)** | AGCAAACAGTAGCTTCCCTG |

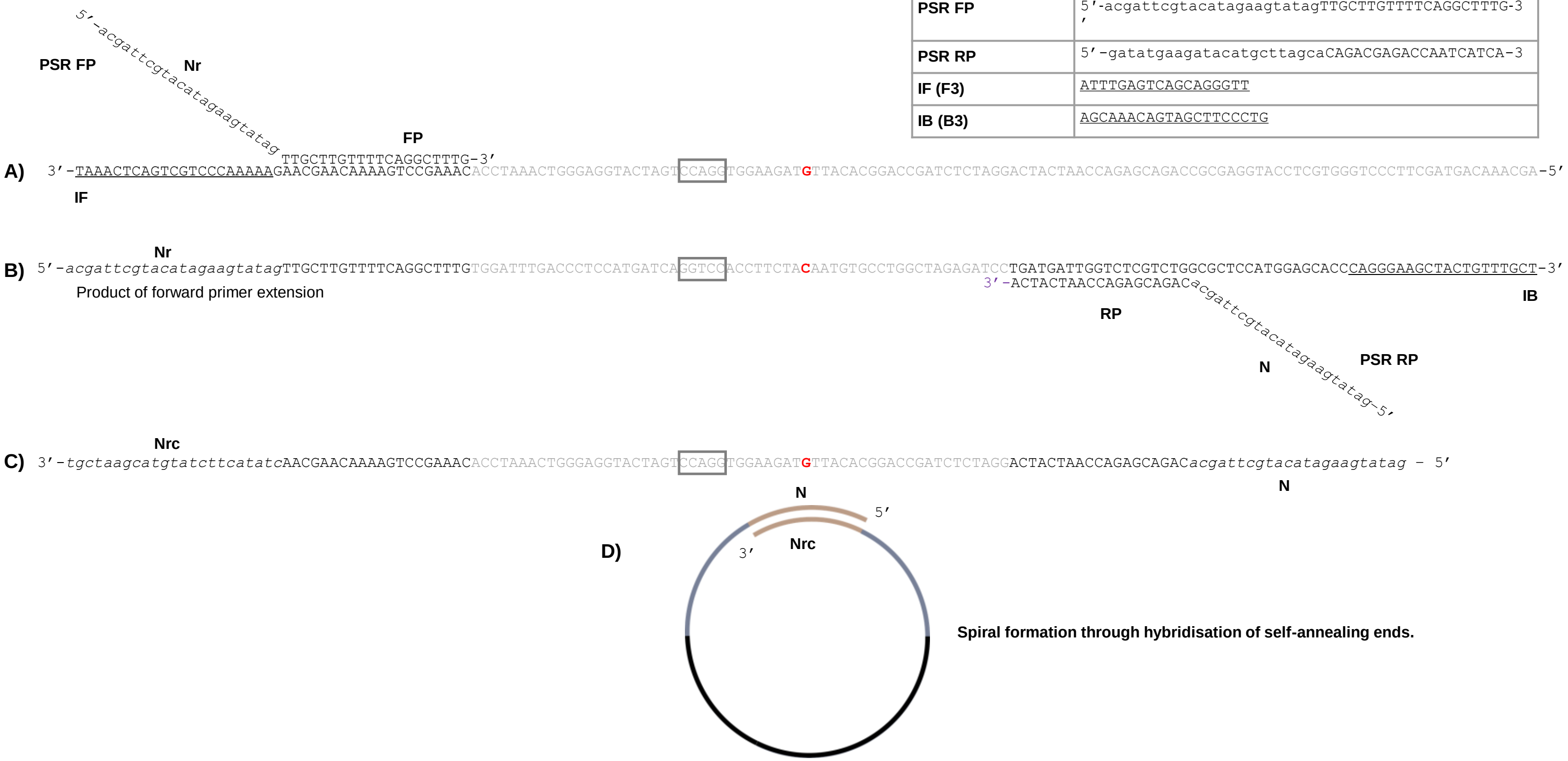


**Supplementary Figure 3. Forward primer ’FP’ initiation of PSR. A)** The 3’-end of the FP anneals to and extends *ESR1* template DNA. New strand synthesis is subsequently displaced by the action of outer IF primer. **B)** The 3’-end of the RP is then able to anneal to and extend this product strand, which is again displaced by the action of an outer strand displacement primer (IB). The resulting product strand **C)** now incorporates reverse complementary (‘rc’) sequences N and N’rc’ of botanic origin, facilitating self-priming from the 3’-open end, spiral formation and exponential amplification **D)** Reverse primer reaction initiation (not shown) would be occurring simultaneously, yielding spiral extension of another single-stranded chain from the 3’-open end. In this case, FP and RP flank but are not selective for the E380Q mutation (single base responsible for mutation shown in red). A typical positive PSR produces a ladder of multiple bands (spirals) by gel electrophoresis, which can be diagnostically digested down to single spiral size (96 bp) using an appropriately selected single site digest enzyme (*AvaII*, grey box).

**Supplementary Table 1: Selective E380Q Mini-LAMP limit of detection.** Data from an average of 2 independent experiments each performed in triplicate, presented as mean TTP ± S.E.M. Limit of detection indicated by an asterisk (*) maintaining a time delay >10 minutes ahead of non-selective NTC amplification.

| **Template copies:** | **Mutant template TTP (min ± SEM)** | **WT template TTP (min ± SEM)** | **Delay (min) between mutant and WT template TTP** |
|---|---|---|---|
| $1 \times 10^7$ | 30.6 ± 0.3 | 42.2 ± 0.6 | 11.6 |
| $1 \times 10^6$ | 35.9 ± 0.7 | 48.2 ± 0.5 | 12.3 |
| $1 \times 10^5$ | 46.0 ± 0.7 | 58.7 ± 1.2 | 12.7 |
| $1 \times 10^4$ | 53.5 ± 1.4 | 70.9 ± 2.9 | 17.4 |
| $1 \times 10^3$ * | 61.8 ± 0.8 | 72.3 ± 4.5 | 10.5 |
| $1 \times 10^2$ | 73.2 ± 3.1 | 73.6 ± 4.1 | 0.4 |
| $1 \times 10^1$ | 78.9 ± 5.9 | 78.3 ± 7.5 | - 0.6 |
| NTC | 81.5 ± 7.5 | 83.6 ± 5.1 | - |

**Supplementary Table 2. Mini-LAMP E*SR1* E380Q selective amplification within a mixed WT and mutant template population.** Mixed ratio mutant and WT synthetic DNA (totalling 1 x $10^4$ copies per reaction) were prepared at 100:0, 75:25, 50:50, 25:75, 0:100, plus NTC. Two individual experiments were performed in quadruplicate. Time to positive (TTP) is the time (minutes) at which amplification exceeds the manually set, reaction consistent threshold and enters the rapid linear, exponential phase.

| Mutant : WT (%) 1 x $10e^4$ copies total: | TTP (min ± SEM) | Delay (min) between mutant and WT template TTP |
|---|---|---|
| **100 : 0** | 56.5 ± 1.9 | - |
| **75 : 25** | 58.7 ± 1.5 | 14.9 |
| **50 : 50** | 60.0 ± 0.86 | 12.7 |
| **25 : 75** | 62.2 ± 1.4 | 11.4 |
| **0 : 100** | 71.4 ± 2.6 | 9.2 |
| **NTC** | 78.0 ± 2.0 | - |
